# Correlated Functional Connectivity and Glucose Metabolism in Brain White Matter Revealed by Simultaneous MRI/PET

**DOI:** 10.1101/2021.04.16.440245

**Authors:** Bin Guo, Fugen Zhou, Muwei Li, John C. Gore, Zhaohua Ding

## Abstract

Blood oxygenation level-dependent (BOLD) signals in white matter (WM) have usually been ignored or undetected, consistent with the lower vascular density and metabolic demands in WM than in gray matter (GM). Despite converging evidence demonstrating the reliable detection of BOLD signals in WM evoked by neural stimulation and in a resting state, few studies have examined the relationship between BOLD functional signals and tissue metabolism in WM. By analyzing simultaneous recordings of MRI and PET data, we found that the correlations between low frequency resting state BOLD signals in WM are spatially correlated with local glucose uptake, which also covaried with the amplitude of spontaneous low frequency fluctuations in BOLD signals. These results provide further evidence that BOLD signals in WM reflect variations in metabolic demand associated with neural activity, and suggest they should be incorporated into more complete models of brain function.

## Introduction

Since the discovery of correlations between spontaneous low frequency fluctuations in blood oxygenation level dependent (BOLD) signals around the mid-90s (Biswal et al. 1995), resting state functional connectivity (FC) has been extensively studied, leading to the identification of several resting state networks in the human brain (Fox et al. 2007). Whilst the vast majority of these studies have hitherto focused on cortical gray matter (GM), there has been growing interest in the evaluation of functional networks in white matter (WM) (Peer et al. 2017; Li et al. 2019). In particular, it has been recently demonstrated that spontaneous low frequency fluctuations in WM BOLD signals are robustly detectable and reflect specific neural activities (Ding et al. 2013; Ding et al. 2018), which suggests the potential of analyzing and characterizing FC in WM.

Notwithstanding compelling evidence provided by experimental studies (Gore et al. 2019) and supportive clinical data that have recently emerged (Faragó et al. 2019; Gao et al. 2019; Wang et al. 2019; Frizzell et al. 2020; Zhang et al. 2020; Cui et al. 2021; Liu et al. 2021; Sarma et al. 2021;), the interpretation of the observed fluctuations in WM signals remains unclear (Gawryluk et al. 2014). Physiologically, the vascular density of WM is approximately one fourth that of GM, so hemodynamic responses to increases in energy demand in WM are expected to be proportionally reduced, so that BOLD signals are weaker and may fall below the sensitivity of conventional acquisitions and analyses. Moreover, it is not clear what processes within WM modulate local metabolic needs or regulate flow and oxygenation to couple neural activity and vascular hemodynamics. The observed signal fluctuations in WM could plausibly originate from venous draining effects from upstream GM or other non-neural confounds that impact BOLD signals throughout the brain parenchyma.

Although previous studies have shown WM BOLD signals are affected concomitantly with changes in neural activity in cortex, there is a residual need to clarify whether those changes reflect an intrinsic metabolic demand within WM itself. The primary energy substrate of brain tissues is glucose, and variations in glucose uptake reflect variations in baseline metabolic rates. We hypothesized the engagement of regions of WM in brain functions in a resting state is reflected in the magnitudes of the spontaneous fluctuations in local BOLD signals and in the strengths of the correlations of BOLD signals across time with other areas, which is interpreted as functional connectivity (FC). We analyzed PET and MRI data previously acquired and reported by (Jamadar et al. 2020). We demonstrate that, by analyzing simultaneous recordings of the uptake of fluorodeoxyglucose (FDG) by dynamic positron emission tomography (PET) (Lameka et al. 2016) and BOLD signals by functional magnetic resonance imaging (fMRI), there are strong and significant spatial correlations between FDG uptakes and FC in WM, and FC is associated with the fractional amplitude of low frequency fluctuations (FALFF) in the BOLD signals. These observations lend strong support to the notion that BOLD signal fluctuations in WM are linked to neural activities through local variations in aerobic metabolism.

## Results

### Measurements of FC, FDG uptake and FALFF

FC, FDG uptake and FALFF measures were computed on the basis of individual WM bundles, which here are denoted as bFC, bFDG and bFALFF. Average bFC, bFDG and bFALFF values across all the bundles and subjects studied and all six sessions ranged from −0.06 to 0.47 (0.27±0.11), 0.47 to 1.00 (0.72±0.15), and 0.20 to 0.25 (0.23±0.01) respectively. Spatial distributions of bFC and bFDG (see left and middle panels in Figure 1A) were highly symmetrical, with a Pearson’s correlation coefficient of 0.966 and 0.995 respectively comparing the bilateral WM bundles. By comparison, the inter-hemispheric similarity of bFALFF distributions (see the right panel in Figure 1A) was somewhat reduced (r=0.744).

**Figure 1.**
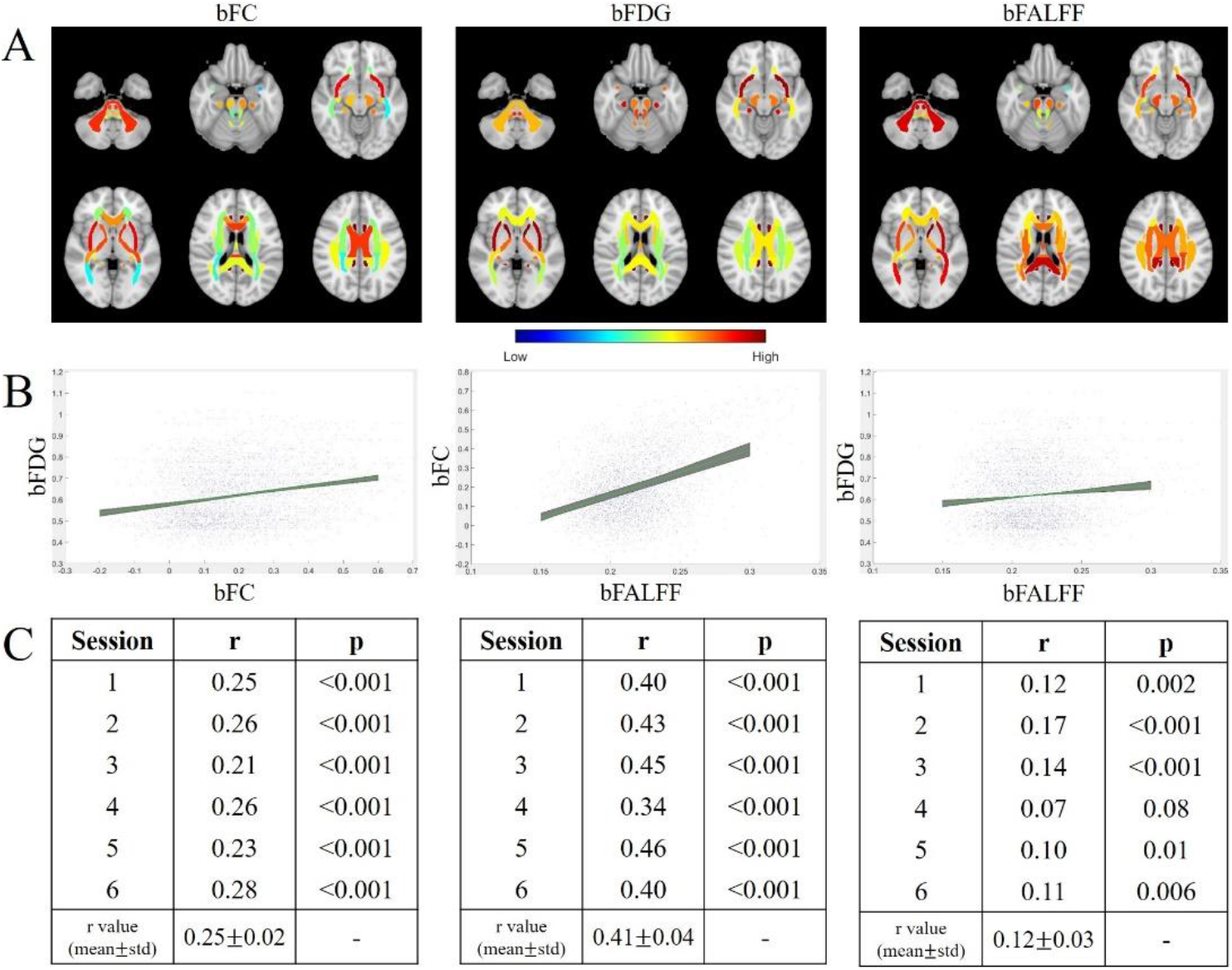
Distributions of bFC, bFDG and bFALFF measures and pairwise correlations between them. (A) Distributions of average bFC (left), bFDG (middle), and bFALFF (right) in selected axial slices (see Figure S1-S3 for full brain distributions). Note that for visualization purposes, average values of these measures are mapped to the original atlas with no WM mask erosions. (B) Scatter plots of linear relationships between bFC and bFDG, bFALFF and bFC, and bFALFF and bFDG. Tight regions shaded in green illustrate highly consistent linear fitting of the three measure pairs across six fMRI sessions. (C) Summary of Pearson’s correlations between the three measure pairs for each of the six fMRI sessions.

### Correlations between bFC and bFDG

Pearson’s correlations between bFC and bFDG pooled over all the WM bundles and subjects studied are shown in Figure 1B (left). As seen, bFC exhibited significant correlation with bFDG, which was highly consistent across the six fMRI sessions (r=0.25±0.02, all p<0.001). bFC was also found to correlate significantly with bFALFF for each session (all p<0.001, r=0.41±0.04) (Figure 1B middle), likely because measures of FC are larger when FALFFincreases compared to physiological noise. For completeness, Pearson’s correlation between bFALFF and bFDG was also computed (Figure 1B right), which appeared to be weaker and had only five of the six sessions reaching significance (r=0.12 ±0.03, all p<0.01 except for session 4).

To assess the amount of variance in a dependent variable explainable by a predictor variable in the absence of intra-subject variations, a mixed effects model was evaluated by treating the subject number as a random effect in each of the three paired comparisons. The p-values for all the three models were <0.001, and the R^2^-Adjusted = 17%, 42% and 8% respectively for bFC vs. bFDG, bFALFF vs. bFC and bFALFF vs. bFDG correlations (see Table 1). The corresponding coefficients for the three pairs of correlations were 0.41, 0.65 and 0.28 respectively, indicating the existence of moderate to strong correlations between these measures when the random effects introduced by inter-subject variations were controlled.

**Table 1.**
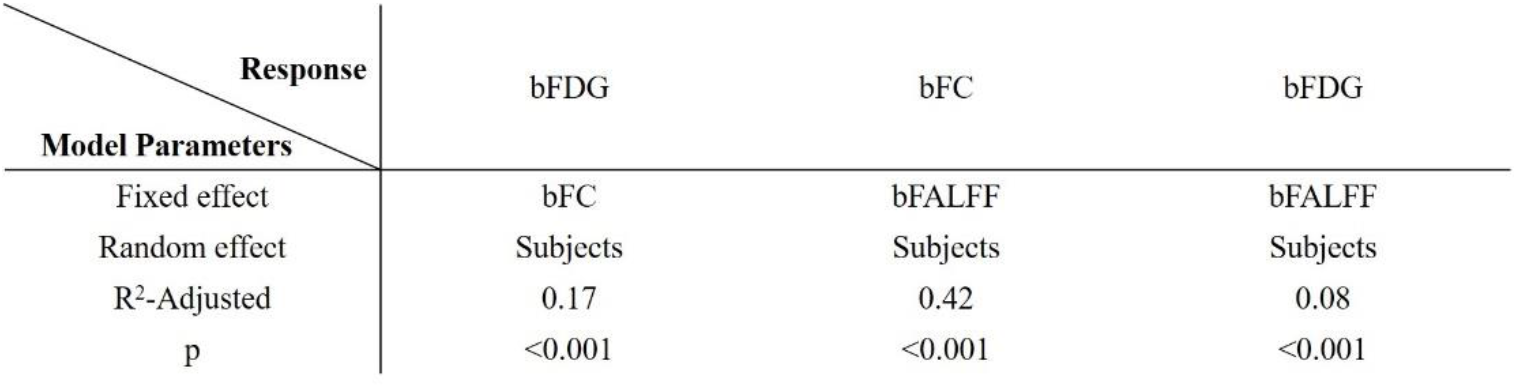
Summary statistics of mixed effect models. See text for explanations.

### Assessments of partial volume effects in WM

To examine whether the observed correlation between bFC and bFDG in WM was corrupted by the effects of partial volume averaging with GM, the correlation derived with WM masks eroded from 0 to 4 mm was compared (see Figure 2). It can be seen that, with the level of erosions increasing from 0 to 3 mm, the correlation varied from r=0.37±0.03 to 0.25±0.02. However, the correlation tended to stabilize with further erosions of WM masks. This trend indicated that with WM masks eroded at 3 mm, the effects of partial volume averaging were quite minimal if any. Also note that the gradual decrease in the correlation coefficient from WM mask erosions of 1-3 mm may be due, at least in part, to the density gradient of interstitial neurons in WM, which tend to be more abundant toward the superficial WM zone (Sedmak et al. 2019).

**Figure 2.**
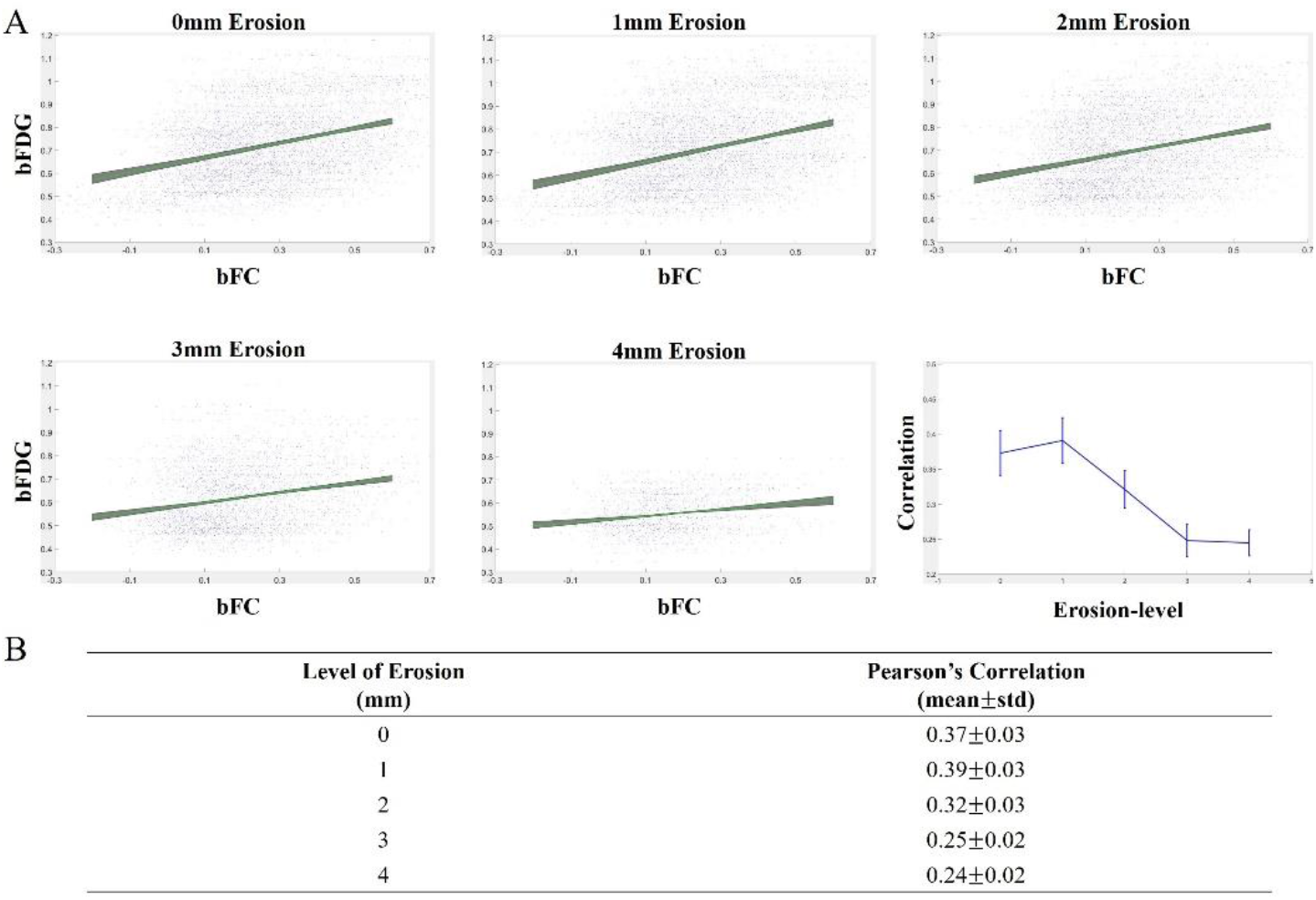
Effects of WM mask erosions on correlations between bFC and bFDG. (A) Linear fittings between bFC and bFDG and correlation coefficients at the levels of WM mask erosions from 0 to 4 mm. (B) Mean and standard deviation of correlation coefficient across six imaging sessions at each level of WM mask erosion. Note that WM masks eroded at 3 mm were used throughout this study.

## Discussion

White matter constitutes nearly half the volume of the human brain, in which axonal fibers serve as information conduits that transmit neural activities between cortical regions. A complete understanding of brain functional architecture therefore requires both GM and WM be taken into consideration. At present, this is only partly achieved by combining functional imaging of GM cortices and structural information from diffusion based tractography of WM pathways (Horn et al. 2014). However, diffusion based tractography does not reveal how WM tracts are engaged in dynamic functional processes within brain networks.

As mentioned previously, there have been several reports of successful detections of functional signals in WM using fMRI (Grajauskas et al. 2019), but their interpretation remains unclear. It has been demonstrated that BOLD signals in GM are correlated with local field potentials (Logothetis 2001), and much of the energy consumption of cortex is accounted for by neural activity of a type not found in WM. This raises concerns as to whether the observed fluctuations in WM reflect BOLD effects consequent on transient variations in metabolic demand. Interestingly, subsequent studies have found that BOLD signals are also correlated with postsynaptic spiking activity, which also increases oxygen consumption (Heeger&Ress, 2002; Mukamel et al. 2005). Direct measurements of metabolism promise to clarify the origins of MRI signal changes. It was found earlier that, although WM has about one-fourth of the vascular density of GM, the oxygen extraction fraction is quite uniform throughout the brain parenchyma (Raichle et al. 2000). A combined PET and fMRI study found that, during a hypocapnia challenge, BOLD signals increased in WM but with reduced magnitudes compared to GM, an effect that was largely attributable to reduced blood flow and volume in WM (Rostrup et al. 2000). Moreover, it was recently observed that global glucose metabolic activities in WM, measured by FDG-PET, vary with the functional state of the brain (Thompson, 2016). These findings suggest that the blood volume and oxygenation level in WM could also fluctuate with neural activities, thereby producing BOLD effects similar but smaller than those found in GM. We therefore analyzed regional glucose metabolism using dynamic PET data and explored their relations with BOLD signals. It was found that FC in WM bundles is significantly correlated with local metabolism and that the FC is associated with the power density of low frequency fluctuations of BOLD signals. These findings support the notion that BOLD signals in WM are modulated by cortical activities and reflect local metabolic variations and are robustly detectable.

It should be noted that fMRI is well known for being susceptible to a variety of artifacts, which contribute high variance to BOLD signals and confound their interpretations (Power et al. 2017). In the context of WM fMRI, a particular concern relates to the possible contributions of vascular drainage from upstream GM (Bianciardi et al. 2011), which adds to other potential confounds common across the brain such as cardiac pulsations, pulmonary modulations, or subject head movements. Our findings that FC in WM bundles is highly and significantly correlated with local FDG uptakes essentially exclude the effects of these confounds as they in principle bear little relations with local metabolisms. There is a possibility that the bulk motion induced by some of these confounds might mediate artificial couplings of FDG and BOLD measurements, but the frequency domain of the bulk motion is presumably different from that of neural activities and after corrections, its impacts can be largely reduced (Behzadi et al. 2007). In light of the correlation between FC and low frequency fluctuations found in WM, it is quite unlikely that the bulk motion contributed significantly, if any, to the observed high correlations between functional connectivity and glucose uptake in WM.

## Methods

A schematic diagram of data analysis for this study is shown in Figure 3.

**Figure 3.**
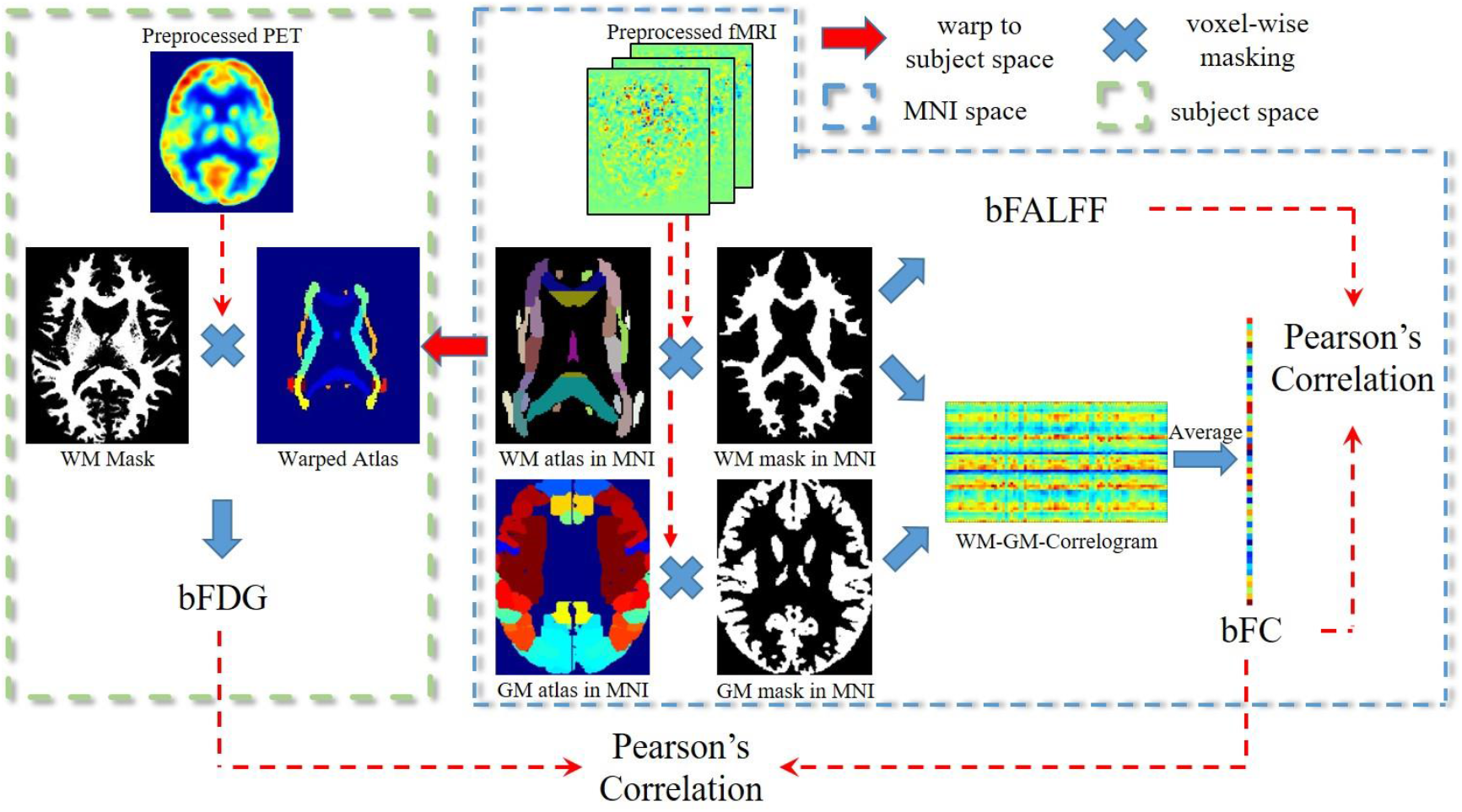
Schematic diagram of analysis framework. See text for detailed explanations of the procedures.

### Participants

The subjects in the current study included 25 right-handed human adults (aged 18-23 yrs, 18 F and 7 M) with no diagnosed mental illness, diabetes or cardiovascular illness. Other inclusion and exclusion criteria are seen in previous reports (Jamadar et al. 2019b, Jamadar et al. 2020),

### MR-PET Imaging

Detailed imaging parameters and procedures can be found elsewhere (Jamadar et al. 2019b). Briefly, each participant underwent a 95-min simultaneous MRI-PET scan in a supine position in a Siemens (Erlangen) 3T Biograph molecular MR (mMR) scanner (Syngo VB20 P). The infusion of [18F] FDG (36 mL/hr) was synchronized with the start of PET acquisitions. In the initial 30 mins, while the PET signal rose to a detectable level, only non-functional MRI scans were acquired, including T1 3D MPRAGE, and some other scans that are not reported in this study. This procedure was followed by six sessions of resting-state PET-fMRI, wherein each lasted for 10 minutes and subsequently went through a series of processing procedures, as described below.

### Preprocessing of PET

For each individual, motion correction was performed on 225 PET volumes using the realign module in SPM (https://www.fil.ion.ucl.ac.uk/spm/software/spm12/) where the first volume was regarded as reference. Each of the realigned PET volumes was then coregistered to the T1 weighted image of the same individual. Summing these coregistered volumes produced a static PET image, from which measurements of FDG uptakes were extracted.

### Preprocessing of Resting State fMRI Data

Resting state data preprocessing involved a few steps as follows. First, the fMRI images were corrected for slice timing and head motion. Second, T1 weighted images were segmented into GM, WM and cerebrospinal fluid (CSF) using SPM, and all these images were registered to the fMRI data space of each individual. Third, mean signals from the whole brain mask were regressed out as nuisance covariates from the fMRI time series. Fourth, the fMRI data, along with the coregistered T1 weighted images as well as the GM and WM segments, were normalized into the Montreal Neurological Institute (MNI) space. Fifth, linear trends from the BOLD images were removed to correct for signal drift.

### Computation of FALFF

To compute FALFF, we defined 48 WM bundle templates based on the JHU-ICBM WM atlas. These WM templates were multiplied by the WM segment obtained in the preceding step, which was thresholded at 0.5 and eroded for 3 mm to eliminate potential partial volume effects from GM (see Figure S4). The time-series in each bundle template were averaged, which was then normalized to unit variance and Fourier transformed to derive the power spectrum. The FALFF was defined to be the average of the square root of the low-frequency power spectrum (0.01~0.05Hz) divided by that from the full frequency range, similarly to (Tomasi et al. 2013).

### Computation of FC

Following the procedure in (Ding et al. 2018), a set of time series was extracted from 130 regions of interest (ROI), including 82 Brodmann areas (BAs) in GM and 48 WM bundles (based on the eroded WM mask as above) for each subject. Each time series was temporally filtered using a bandpass filter (0.01~0.1Hz). Pearson’s correlation in the time series was calculated for each pair of WM and GM regions. This resulted in an 82*48 FC matrix, from which BA-averaged FC was computed for each of the WM bundles that represented its overall FC profile.

### Computation of FDG uptake

The FDG uptake was computed by first normalizing the image intensity of each static PET voxel with the global mean of the entire brain, so that the mean value of the entire brain is 1. Then WM bundle templates defined above were warped back to each individual space of the PET data, from which averaged FDG uptake values from each bundle was obtained.

## Acknowledgments

This work was supported by the National Institutes of Health (NIH) grants R01 NS093669 (J.C.G) and R01 NS113832 (J.C.G), the National Key R&D Program of China Grants 2018YFA0704100 and 2018YFA0704101, and the National Natural Science Foundation of China Grant 61601012. We wish to sincerely thank Dr. Hakmook Kang (Department of Biostatistics, Vanderbilt University School of Medicine) for his valuable advice on our statistical analysis.

