## supplementary material for "Correlated Functional Connectivity and Glucose Metabolism in Brain White Matter Revealed by Simultaneous MRI/PET"

### Supplementary Materials

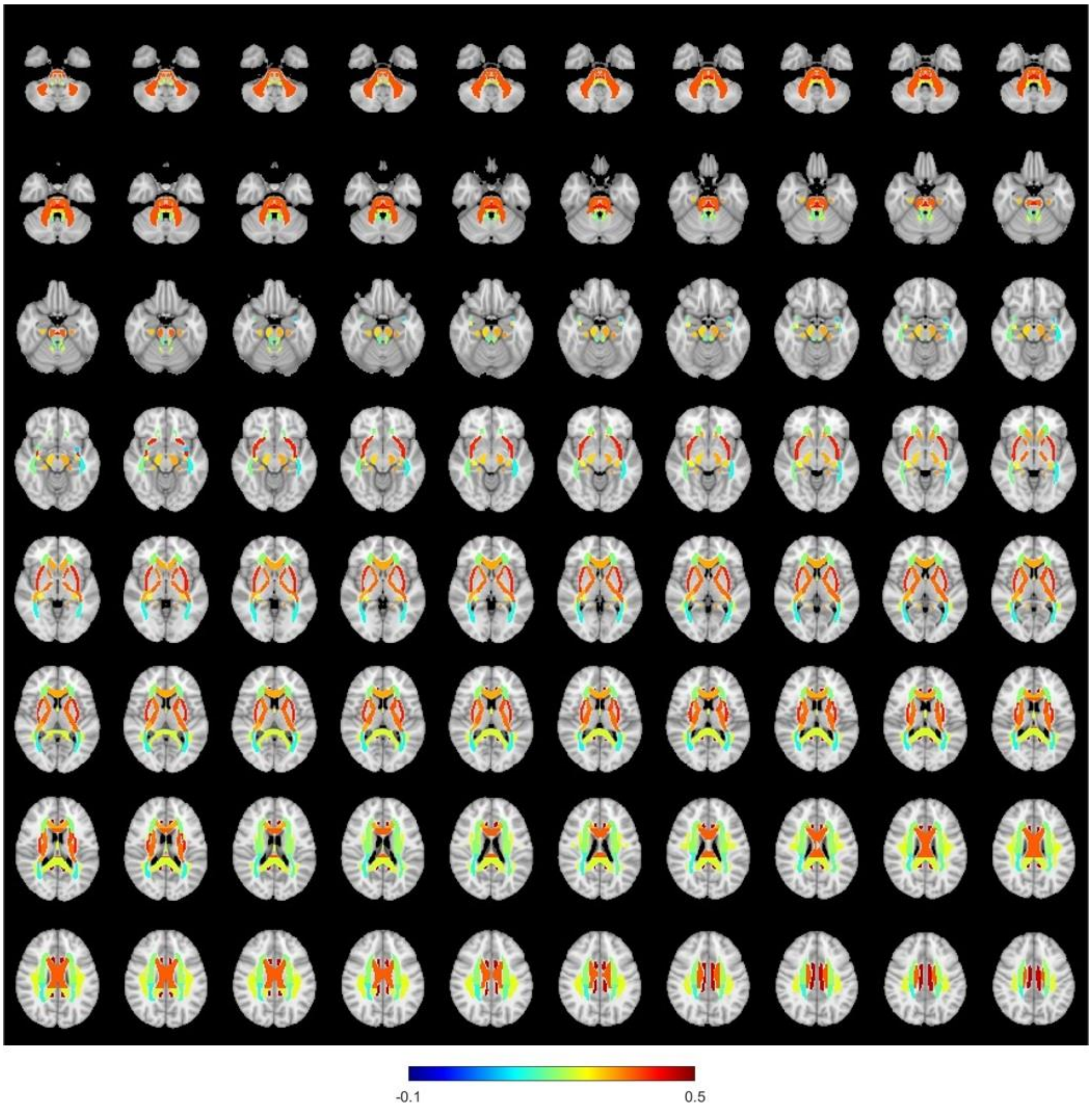


Figure S1. Distributions of bFC. Bundle values are averaged across all the subjects studied and all six imaging sessions, and are superimposed onto the anatomical images in the MNI space.

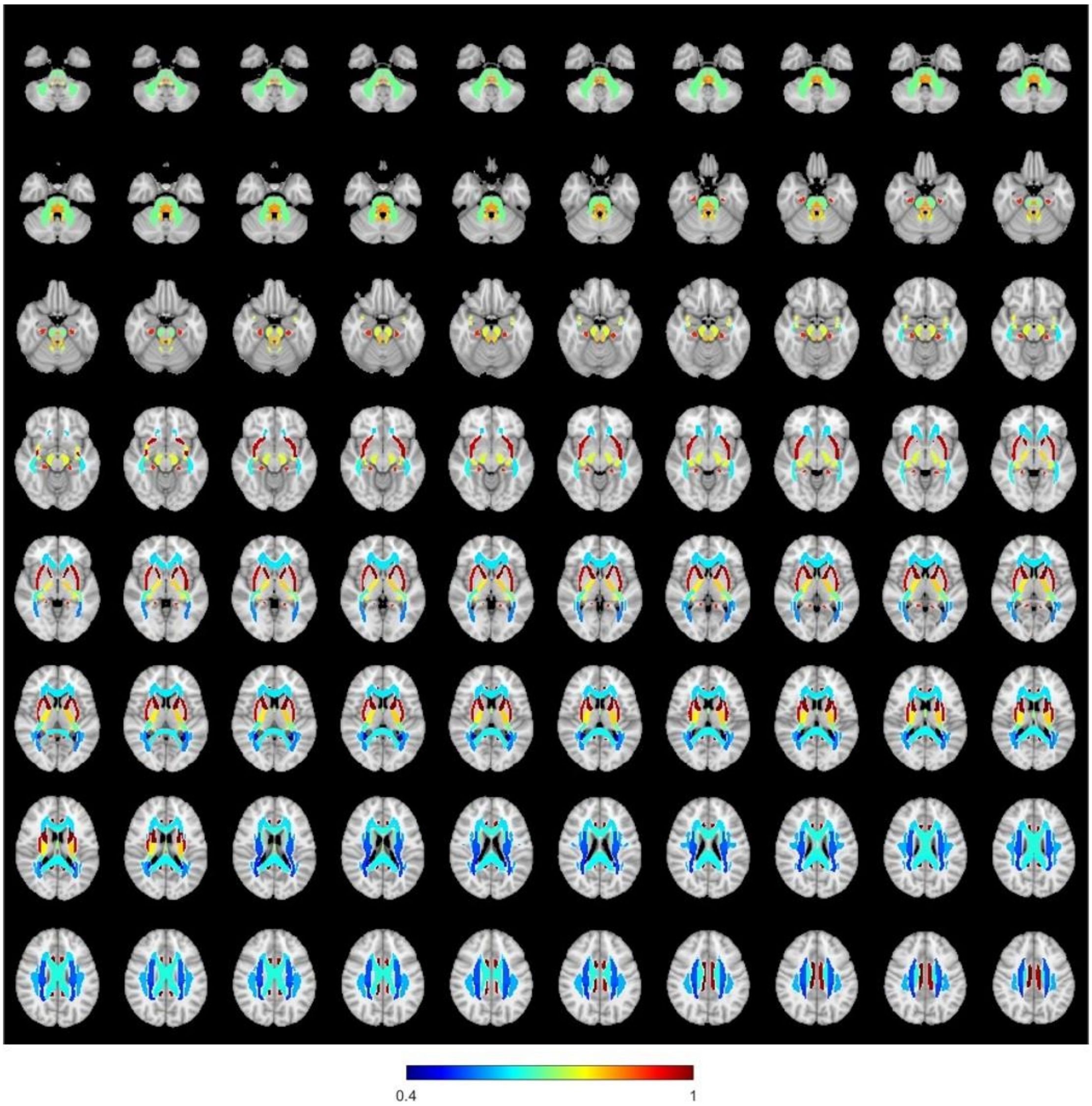


Figure S2. Distributions of bFDG. Bundle values are averaged across all the subjects studied and all six imaging sessions, and are superimposed onto the anatomical images in the MNI space.

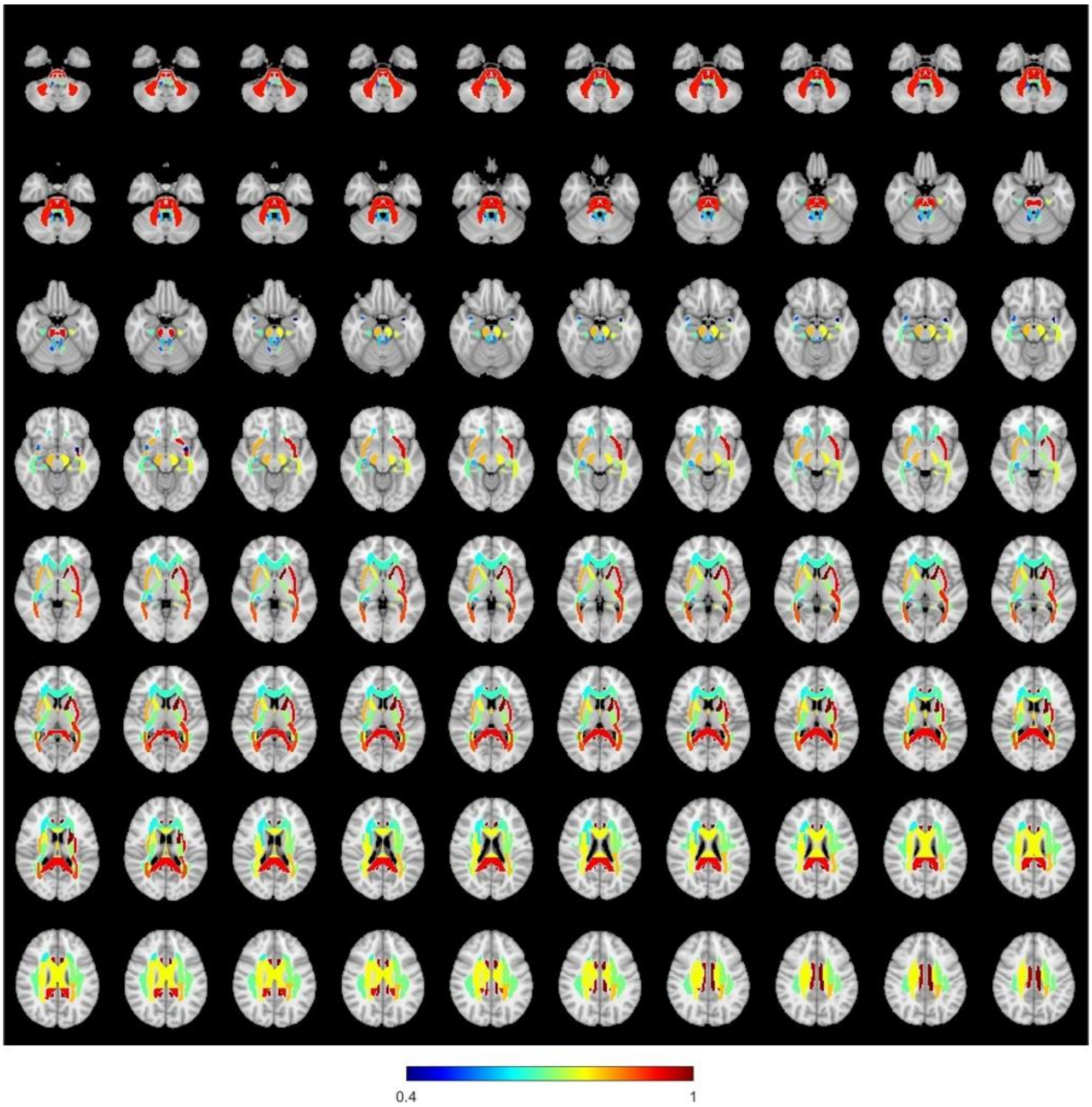


Figure S3. Distributions of bFALFF. Bundle values are averaged across all the subjects studied and all six imaging sessions, and are superimposed onto the anatomical images in the MNI space. Note that the values of bFALFF have been rescaled from the original range of [0.20, 0.25] to [0.40, 1] for enhanced visualization.

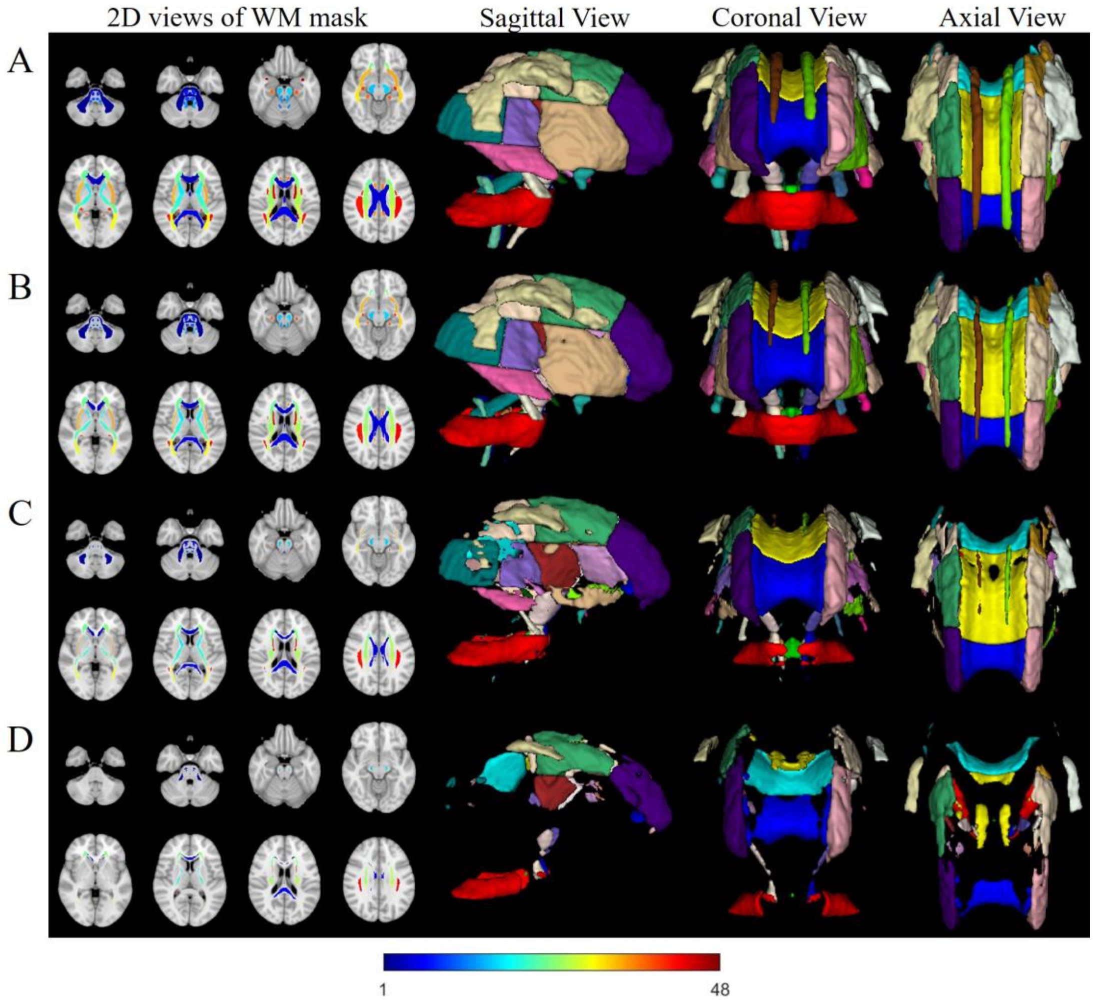


Figure S4. Illustrations of WM masks before erosion (A), after erosion of 1 mm (B), 2 mm (C), and 3 mm (D). 3D renderings are shown in the right three columns for sagittal, coronal and axial views.
